# Neutralizing antibody evasion of SARS-CoV-2 JN.1 derivatives KP.3, KP.3.1.1, LB.1, and XEC

**DOI:** 10.1101/2024.11.04.621772

**Authors:** Kaori Sano, Kei Miyakawa, Hideaki Kato, Yayoi Kimura, Atsushi Goto, Akihide Ryo, Shinji Watanabe, Hideki Hasegawa

**Author notes:** **Co-corresponding Authors: Kei Miyakawa, Ph.D., Shinji Watanabe, D.V.M., Ph.D., Hideki Hasegawa, M.D., Ph.D.**, Research Center for Influenza and Respiratory Viruses, National Institute of Infectious Disease, 4-7-1 Gakuen, Musashimurayama, Tokyo 208-0011, Japan.

## Abstract

The emergence of SARS-CoV-2 variants poses ongoing challenges to vaccine efficacy. We evaluated neutralizing antibody responses against JN.1 and its derivatives (KP.3, KP.3.1.1, LB.1, and XEC) in healthcare workers who received seven doses of BNT162b2, including XBB.1.5 monovalent vaccine. In COVID-19-naïve individuals, KP.3.1.1 and LB.1 showed substantial immune escape, while previously infected individuals maintained neutralization activity against all variants. We also demonstrated that JN.1-based immunization induces robust cross-neutralizing activity against emerging variants. A single amino acid deletion at position 31 in the spike protein significantly impacted immune evasion. These findings support the potential effectiveness of JN.1-based vaccines while highlighting the need for continued surveillance and vaccine optimization.

## Main

The COVID-19 pandemic remains a global health concern, particularly due to the continuous emergence of SARS-CoV-2 variants that potentially compromise vaccine efficacy, leading to persistent viral infection and post COVID-19 condition (Long COVID) ^1,2^. Recent surveillance data indicate the global spread of the JN.1 subvariant and its derivatives (KP.3, KP.3.1.1, LB.1, and XEC) (**Figure 1A, Supplementary Figure 1**). While JN.1 has been selected as the vaccine strain for next season ^3^, understanding the cross-protective efficacy of the monovalent XBB.1.5 vaccine given last season against these emerging variants remains crucial, especially for vulnerable populations.

**Figure 1.**
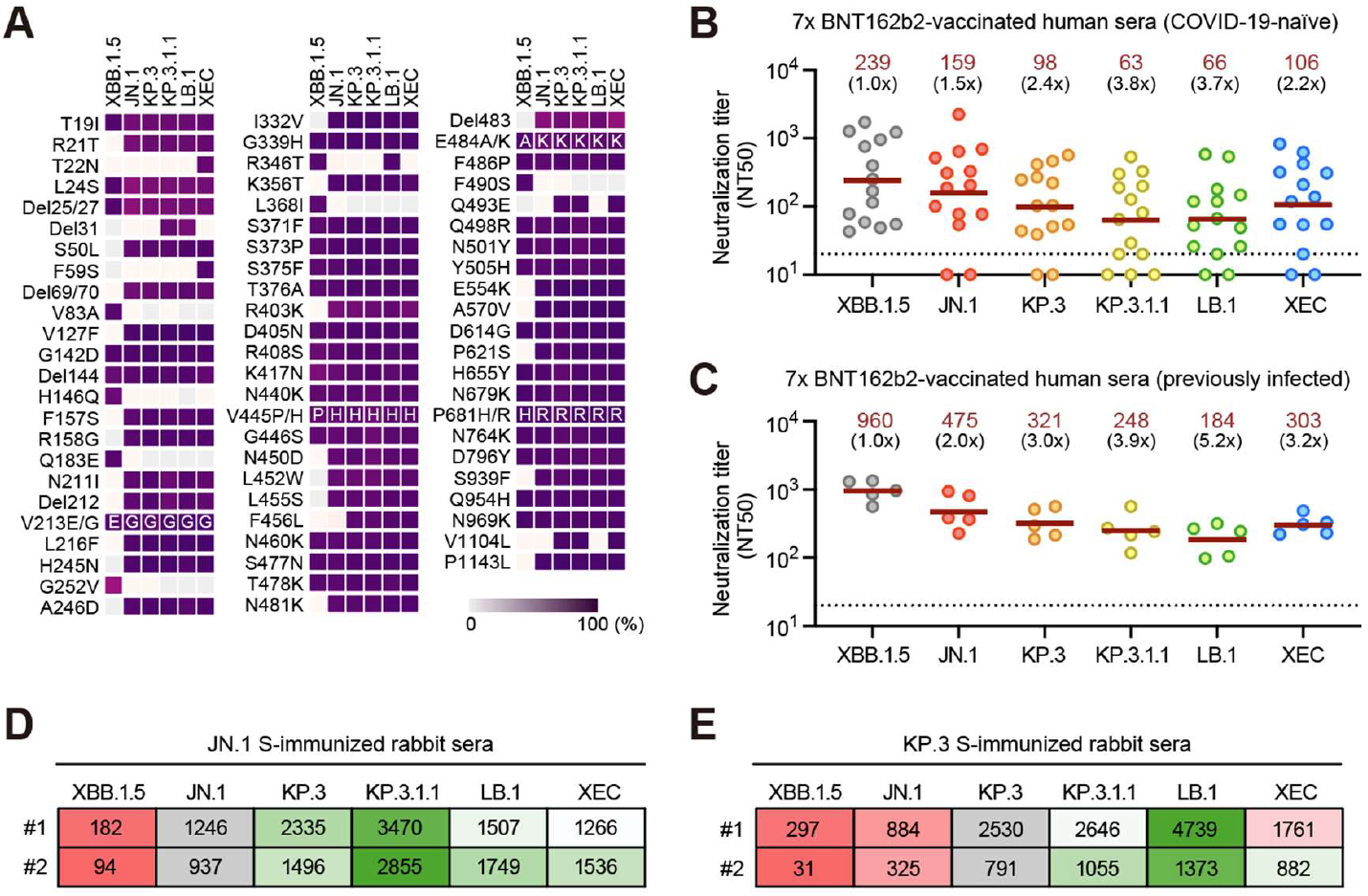
Neutralizing antibody responses to the SARS-CoV-2 JN.1 derivatives. **(A)** Schematic representation of spike (S) protein mutations across SARS-CoV-2 Omicron subvariants. The intensity of the square coloring correlates with mutation frequency. Data were obtained from Outbreak.info (https://outbreak.info/). **(B, C)** Neutralizing antibody titers (NT50) against Omicron subvariants in sera from BNT162b2 vaccinees, stratified by prior SARS-CoV-2 infection status: COVID-19-naïve (**B**, n=14) and previously infected individuals (**C**, n=5). Brown horizontal bars and numbers represent geometric mean titers. Black numbers indicate fold changes in neutralization resistance relative to XBB.1.5 (reference strain). The dotted line indicates the limit of detection for this assay. **(D, E)** Neutralizing antibody titers against Omicron subvariants measured in two rabbit sera immunized with JN.1 (**D**) or KP.3 (**E**) S antigens. Gray cells indicate the immunizing strain (reference). Red and green cells highlight strains showing neutralization resistance and susceptibility, respectively.

We evaluated neutralizing antibody responses in 19 Japanese healthcare workers who received seven doses of BNT162b2 mRNA vaccine, with XBB.1.5 monovalent vaccine as their final dose in December 2023. The cohort included 14 COVID-19-naïve and 5 previously infected individuals (**Supplementary Table 1, 2**). To evaluate the neutralizing antibody responses, pseudovirus neutralization assays ^4,5^ were conducted, with XBB.1.5 as the reference strain. In COVID-19-naïve individuals, when normalized to neutralization titers against XBB.1.5 as baseline, KP.3.1.1 and LB.1 demonstrated substantial immune escape (>3-fold reduction in neutralization), whereas KP.3 and XEC showed moderate evasion (2-to 3-fold reduction) (**Figure 1B**). Similar immune escape patterns were observed in previously infected individuals (**Figure 1C**). Some COVID-19-naïve subjects exhibited undetectable neutralizing activity against JN.1 and its derivatives, while previously infected individuals maintained neutralizing responses against all tested variants (**Figure 1B, C**). Notably, both KP.3.1.1 and LB.1 harbor a deletion at position 31 in the spike (S) protein (**Figure 1A**), highlighting the potential role of this deletion in enhanced immune evasion.

To further characterize variant-specific immune responses, we analyzed cross-reactivity using rabbit antisera against JN.1 and KP.3 S proteins. JN.1 S-immunized sera showed comparable neutralization of KP.3, LB.1, and XEC, with enhanced neutralization activity against KP.3.1.1 (approximately 3-fold increase) (**Figure 1D**). KP.3 S-immunized sera also demonstrated robust neutralization of JN.1 derivatives, although it showed reduced activity against JN.1 itself (2.5-fold decrease) (**Figure 1E**). Sera immunized with JN.1 S or KP.3 S no longer showed robust neutralizing activity against XBB.1.5 (**Figure 1D, E**). These findings reveal that while JN.1 and KP.3 share antigenic properties with currently circulating JN.1 derivatives, they display antigenic differences from each other and from XBB.1.5.

This study demonstrates the robust cross-neutralizing activity of JN.1-based vaccination against emerging variants, including the immune-evading KP.3.1.1 and LB.1, highlighting the potential efficacy of JN.1-based vaccines. However, the small sample size may limit the statistical power and generalizability of the results. Larger studies are needed to confirm these results. Nevertheless, future vaccine development may consider structural modifications, particularly around the 31st amino acid position of the S protein, to improve broad-spectrum protection against emerging variants.

## Acknowledgements

We thank Kenji Yoshihara, Saori Takumi, and Nanako Amano for technical assistance. We also thank Kazuo Horikawa and Takayuki Kurosawa for collecting specimens. This study was supported by grants from AMED (JP24wm0325061 to KS; JP22fk0108518 to KM; and, JP21nf0101626, JP243fa827012, JP243fa827016, and JP243fa727002 to HH) and grants from JSPS (JP23K27419 to KM).

