## Supplemental information for "Neutralizing antibody evasion of SARS-CoV-2 JN.1 derivatives KP.3, KP.3.1.1, LB.1, and XEC"

**Supplementary Figure 1**

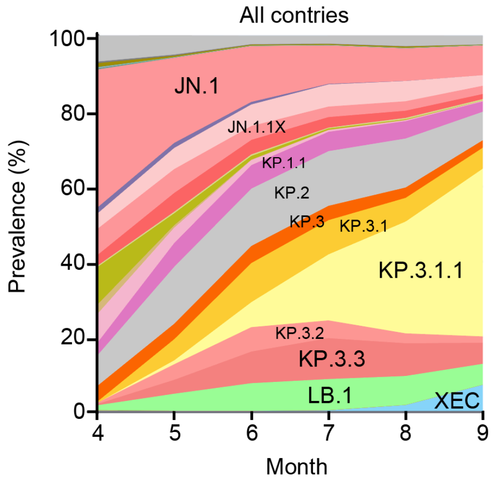


**Supplementary Figure 1. Global outbreak of SARS-CoV-2 variants.**

The prevalence of SARS-CoV-2 variants globally from April 2024 to September 2024. Data were collected from GISAID Lineage Frequency.

**Supplementary Table 1 Study participants (COVID-19-naïve individuals)**

| ID | Age | Sex | Number of vaccinations | Vaccination history* | Vaccination date (YYYY/MM/DD) | | | | | | |
|---|---|---|---|---|---|---|---|---|---|---|---|
| | | | | | 1st | 2nd | 3rd | 4th | 5th | 6th | 7th |
| **001** | 57 | Female | 7 | W/W/W/W/B/B/X | 2021/3/17 | 2021/4/7 | 2021/12/22 | 2022/7/29 | 2022/11/22 | 2023/6/16 | 2023/12/8 |
| **008** | 49 | Female | 7 | W/W/W/W/B/B/X | 2021/3/19 | 2021/4/9 | 2021/12/22 | 2022/8/10 | 2022/11/22 | 2023/6/8 | 2023/11/22 |
| **009** | 45 | Female | 7 | W/W/W/W/B/B/X | 2021/3/23 | 2021/4/13 | 2021/12/20 | 2022/7/31 | 2022/11/28 | 2023/6/14 | 2023/12/1 |
| **012** | 61 | Male | 7 | W/W/W/W/B/B/X | 2021/3/22 | 2021/4/12 | 2021/12/20 | 2022/7/26 | 2022/11/28 | 2023/6/14 | 2023/11/22 |
| **019** | 48 | Female | 7 | W/W/W/W/B/B/X | 2021/3/15 | 2021/4/5 | 2021/12/21 | 2022/7/23 | 2022/11/22 | 2023/6/16 | 2023/12/8 |
| **020** | 52 | Female | 7 | W/W/W/W/B/B/X | 2021/3/19 | 2021/4/9 | 2021/12/22 | 2022/8/5 | 2022/12/23 | 2023/6/16 | 2023/12/15 |
| **021** | 56 | Male | 7 | W/W/W/W/B/B/X | 2021/3/18 | 2021/4/8 | 2021/12/21 | 2022/7/26 | 2022/12/7 | 2023/6/7 | 2023/11/22 |
| **023** | 34 | Male | 7 | W/W/W/W/B/B/X | 2021/3/17 | 2021/4/7 | 2020/12/20 | 2022/7/30 | 2022/12/9 | 2023/6/9 | 2023/12/15 |
| **026** | 50 | Male | 7 | W/W/W/W/B/B/X | 2021/3/15 | 2021/4/5 | 2021/12/20 | 2022/7/14 | 2022/11/5 | 2023/6/20 | 2023/12/15 |
| **029** | 42 | Male | 7 | W/W/W/W/B/B/X | 2021/3/18 | 2021/4/8 | 2021/12/24 | 2022/8/6 | 2022/12/9 | 2023/6/15 | 2023/12/1 |
| **030** | 43 | Female | 7 | W/W/W/W/B/B/X | 2021/3/15 | 2021/4/5 | 2021/12/23 | 2022/8/2 | 2022/12/9 | 2023/6/20 | 2023/12/1 |
| **036** | 39 | Female | 7 | W/W/W/W/B/B/X | 2021/3/12 | 2021/4/5 | 2021/12/20 | 2022/7/15 | 2022/11/5 | 2023/6/20 | 2023/12/15 |
| **039** | 48 | Female | 7 | W/W/W/W/B/B/X | 2021/3/19 | 2021/4/9 | 2021/12/23 | 2022/8/6 | 2022/11/25 | 2023/6/8 | 2023/12/15 |
| **042** | 39 | Female | 7 | W/W/W/W/B/B/X | 2021/3/19 | 2021/4/9 | 2022/1/24 | 2022/7/27 | 2022/11/23 | 2023/6/16 | 2023/11/22 |

*W, Wuhan monovalent; B, Wuhan/BA bivalent; X, XBB monovalent

**Supplementary Table 2 Study participants (previously infected individuals)**

| ID | Age | Sex | Number of vaccinations | Vaccination history* | Vaccination date (YYYY/MM/DD) | | | | | | |
|---|---|---|---|---|---|---|---|---|---|---|---|
| | | | | | 1st | 2nd | 3rd | 4th | 5th | 6th | 7th |
| **014** | 57 | Female | 7 | W/W/W/W/B/B/X | 2021/3/19 | 2021/4/9 | 2021/12/24 | 2022/7/20 | 2022/12/9 | 2023/6/9 | 2023/12/8 |
| **018** | 49 | Female | 7 | W/W/W/W/B/B/X | 2021/3/22 | 2021/4/12 | 2021/12/21 | 2022/7/30 | 2022/11/22 | 2023/6/9 | 2023/12/1 |
| **022** | 45 | Female | 7 | W/W/W/W/B/B/X | 2021/3/15 | 2021/4/5 | 2021/12/23 | 2022/7/22 | 2022/12/9 | 2023/6/9 | 2023/12/1 |
| **035** | 61 | Male | 7 | W/W/W/W/B/B/X | 2021/3/23 | 2021/4/13 | 2021/12/24 | 2022/7/30 | 2022/11/25 | 2023/6/19 | 2023/12/8 |
| **041** | 48 | Female | 7 | W/W/W/W/B/B/X | 2021/3/17 | 2021/4/7 | 2021/12/22 | 2022/7/22 | 2022/11/2 | 2023/6/20 | 2023/11/22 |

*W, Wuhan monovalent; B, Wuhan/BA bivalent; X, XBB monovalent

### Materials and methods

#### Participants

Study participants were recruited from healthcare workers at the Yokohama City University Hospital, Japan, who had completed seven doses of BNT162b2 mRNA vaccine, with the XBB.1.5 monovalent vaccine administered as the final dose in December 2023. The mean age of the participants was 48.6 years (range: 34–61), and 31.5% were male. Prior to the experiments, all samples were screened for IgG antibodies against the SARS-CoV-2 nucleocapsid (N) protein to determine their serostatus. Blinding was not implemented as the experimental outcomes did not involve subjective assessments. Sample size calculations were not performed for this study. The study protocol was approved by the Yokohama City University Certified Institutional Review Board (approval number: F231100032). Written informed consent was obtained from all participants. This study was conducted according to the principles of the Declaration of Helsinki.

#### Pseudovirus neutralization assay

The neutralization assay was performed as previously described [1,2]. In brief, VeroE6/TMPRSS2/LgBiT cells were inoculated with HiBiT-tagged pseudovirus mixed with serially diluted serum samples. Intracellular luciferase activity was measured 4 hours after inoculation. The 50% neutralization titer (NT50) was defined as the serum dilution factor that resulted in a 50% reduction in luminescence compared to the non-serum control. For samples showing no detectable neutralizing activity to interpolate NT50, a value of 10 was assigned.

#### Recombinant proteins

Recombinant SARS-CoV2 spike (S) proteins were produced using a mammalian cell protein expression system as previously described [3]. DNA fragments encoding the SARS-CoV-2 S gene sequence (GenBank: MN908947) containing strain-specific mutations for JN.1 or KP.3, along with six proline substitutions (Hexapro) [4], were synthesized by Eurofins Genomics. These fragments were cloned into a pCAGGS mammalian expression vector containing a fibritin trimerization domain, strep-tag, and his-tag sequences at the 3' end of the S protein insertion site. Protein expression was performed using the Expi293 Expression System (Thermo Fisher Scientific) following the manufacturer's instructions. The cell culture supernatants were clarified by centrifugation at 4,000 × g, filtered, and purified using Ni-NTA Agarose (QIAGEN). The purified proteins were concentrated using Amicon Ultracell (Merck) centrifugation units, and the buffer was changed to PBS (pH 7.4). The proteins were stored at -80 °C until use.

**Polyclonal antibody production**

Female Slc:JW/CSK rabbits (12 weeks old, n=2 per group) were used to generate polyclonal antibodies against the SARS-CoV-2 spike protein. Pre-immune sera were collected before immunization. The immunization procedures were conducted by Japan SLC, Inc. Briefly, each rabbit received an initial immunization with 1 mg of recombinant SARS-CoV-2 spike protein emulsified with Freund's Complete Adjuvant (Difco Laboratories), followed by a booster immunization 21 days later using the same amount of antigen emulsified with Freund's Incomplete Adjuvant (Difco Laboratories). Immune sera were collected 28 days after the initial immunization. All experimental procedures were performed in accordance with institutional guidelines for animal experimentation.
